# Individual recombinant repeats of MUC16 display variable binding to CA125 antibodies

**DOI:** 10.1101/2023.02.08.527749

**Authors:** Chien-Wei Wang, Eliza K. Hanson, Lisa Minkoff, Rebecca J. Whelan

## Abstract

**BACKGROUND:** Despite its importance in the clinical management of ovarian cancer, the CA125 biomarker—located on the mucin protein MUC16—is still not completely understood. Questions remain about MUC16’s function and structure, specifically the identity and location of the CA125 epitopes.

**OBJECTIVE:** The goal of this study was to characterize the interaction of individual recombinant repeats from the tandem repeat domain of MUC16 with antibodies used in the clinical CA125 II test.

**METHODS:** Using *E. coli* expression, we isolated nine repeats from the putative antigenic domain of CA125. Amino acid composition of recombinant repeats was confirmed by high-resolution mass spectrometry. We characterized the binding of four antibodies—OC125, M11, “OC125-like,” and “M11-like”—to nine recombinant repeats using Western blotting, indirect enzyme-linked immunosorbent assay (ELISA), and localized surface plasmon resonance (SPR) spectroscopy.

**RESULTS:** Each recombinant repeat was recognized by a different combination of CA125 antibodies. OC125 and “OC125-like” antibodies did not bind the same set of recombinant repeats, nor did M11 and “M11-like” antibodies.

**CONCLUSIONS:** Characterization of the interactions between MUC16 recombinant repeats and CA125 antibodies will contribute to ongoing efforts to identify the CA125 epitopes and improve our understanding of this important biomarker.

## 1 Introduction

Considerable effort has been directed toward expanding the suite of biomarkers available for diagnosing and monitoring high-grade serous ovarian cancer (HGSOC) [1 – 8]. Although new biomarkers—most significantly human epididymis protein 4 (HE4) [9, 10]—have been identified and are proving to be transformative, enabling new assays and algorithms [11 – 16], no biomarker has supplanted cancer antigen 125 (CA125), which remains the clinical gold standard for monitoring response to treatment and detecting cancer recurrence [17 – 20]. The FDA-approved assay for CA125 is widely used, despite the fact that the CA125 epitopes have not been defined and controversy persists regarding the minimal functional unit necessary for antibody detection [21 – 23]. In other words, the test that underlies vital decisions in ovarian cancer care employs a mechanism that is not understood.

The CA125 epitopes are carried on MUC16, a large mucin [24, 25]. Some structural features of MUC16 have been determined, including a highly glycosylated N-terminal domain, an immunologically active tandem repeat region containing many similar but non-identical subdomains, and a C-terminal domain including a transmembrane region and cytoplasmic tail [26, 27]. The circulating form of MUC16 detected with the CA125 II ELISA does not contain the transmembrane or cytoplasmic regions. Based on competition studies, CA125 antibodies have been sorted into three groups: OC125-like (group A), M11-like (group B), and OV197-like (group C) [28, 29]. The CA125 II ELISA test is a double determinant immunoassay using two different antibodies (M11 and OC125) as capture and tracer [30]. Earlier iterations of the assay used OC125 as both capture and tracer [31]. The location and identity of the CA125 epitopes remain undefined, and diverse experimental approaches to determine the epitopes have been reported [32 – 36]. Experiments by Bressan and co-workers using Western blot analysis of six recombinantly expressed repeat domains (R2, R7, R9, R11, R25, and R51) revealed that the antibodies used in the CA125 II test (OC125 [31] and M11 [37]) do not recognize all repeat domains uniformly [35]. We hypothesized that variation in antibody recognition may be observed in other recombinant repeats and that the nature of the molecular recognition assay may influence whether binding is observed, particularly if the CA125 epitope is conformational [38].

Here we report the expression of nine recombinant repeats from the putative antigenic domain of CA125 (R2, R5, R6, R7, R9, R11, R25, R34, and R58, in the O’Brien numbering system, sequence alignment shown in SI Figure 1) [26]. Using Western blotting, indirect ELISA, and localized surface plasmon resonance (SPR) spectroscopy, we characterized the interactions of expressed and purified recombinant repeats with four CA125-binding monoclonal antibodies (OC125, M11, “OC125-like,” and “M11-like”). Consistent with our hypothesis and previous reports, the epitopes were found to be distributed nonuniformly over the recombinant repeats, and variation across assay method was observed. Without knowledge of the CA125 epitopes, it is impossible to characterize individual variation in MUC16 proteoforms or to determine whether MUC16 expression changes during cancer development, in response to treatment, or during recurrence. To improve the long-term survival of ovarian cancer patients, there is a vital need to improve the diagnostic value of CA125. This study represents the largest set of MUC16 recombinant repeats, the largest number of antibodies, and the most molecular interaction assay methods reported to date and contributes to ongoing efforts to understand the molecular nature and immunological activity of CA125. The rationale of this study is that its results will ultimately enable us to reinvent the CA125 test by developing new affinity reagents to supplement or replace the antibodies in current use, since a biomarker is only as good as the tools available to detect it.

**Figure 1.**
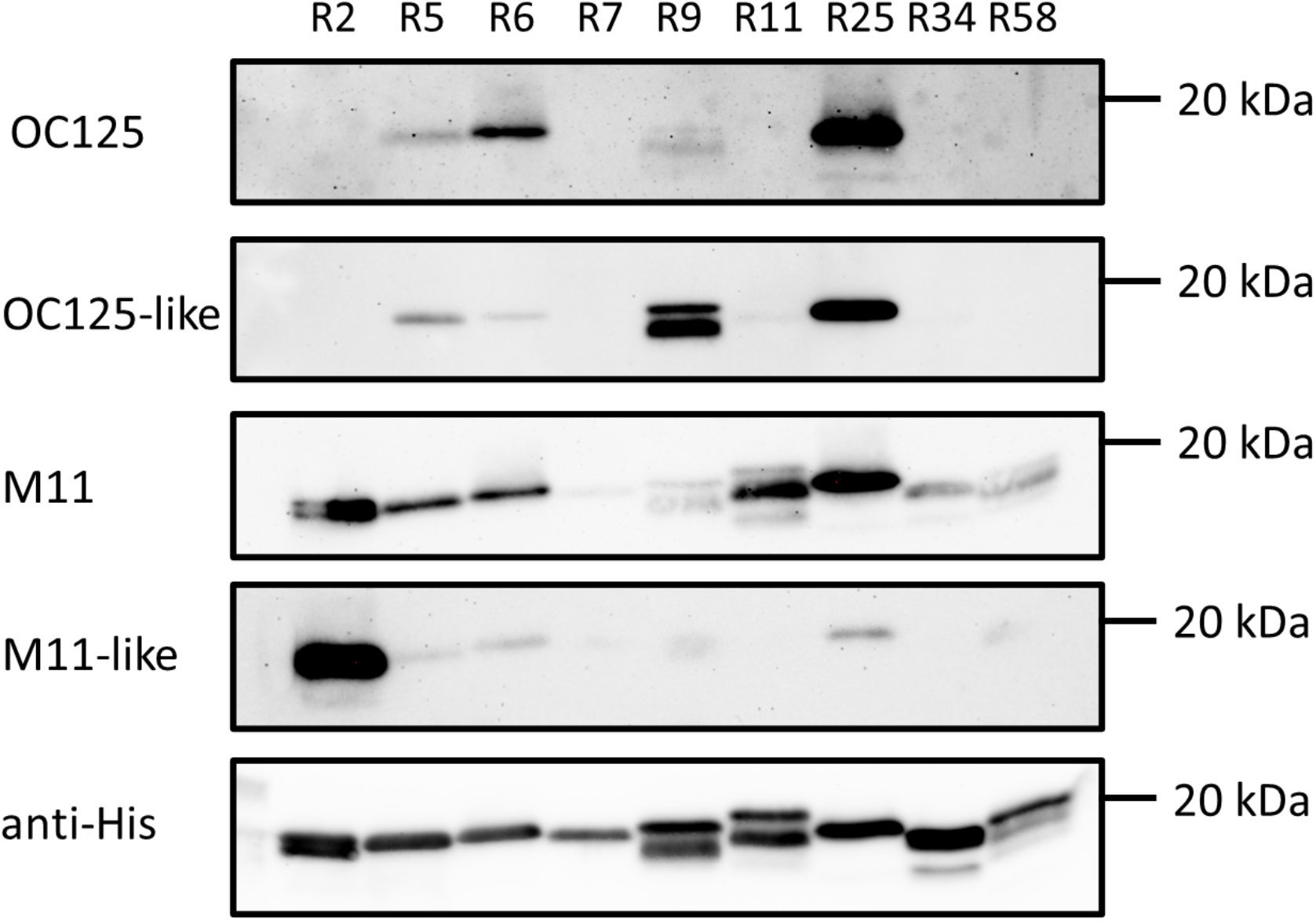
Western blot images of recombinant repeats from MUC16 probed with antibody clones OC125 (1:200), M61704 (OC125-like, 1:2000), M11 (1:100), and M61703 (M11-like, 1:2000), and an anti-His (His.H8, 1:2000) loading control.

## 2 Materials and methods

### 2.1. Recombinant repeat expression and purification

The MUC16 coding sequence as described by O’Brien and co-workers [26] was obtained from NCBI (GenBank: AF414442.2). The nucleotide sequences of nine tandem repeats were synthesized and cloned into pET14b vector (GenScript, Piscataway, NJ), which was used to express protein with N’ 6xHistidine-tagging (6xHis) by XhoI and BamHI sites. Each plasmid was transformed into the SHuffle T7 Express *E. coli* strain (New England Biolabs, Beverly, MA). Bacteria clones were grown in Luria–Bertani (LB) broth (Thermo, Waltham, MA) containing 100 μg/mL ampicillin at 30°C to exponential phase, and protein expression was induced with 400 μM IPTG for 4 hours. Cells were harvested and lysed by freeze-thaw cycles in phosphate buffered saline (PBS) containing cOmpletet™ protease inhibitor (Thermo). The 6xHis-tagged recombinant repeats were then purified on Ni-nitrilotriacetic acid (NTA) beads (Thermo) and verified by Western blot with anti-6xHis antibody and mass spectrometry as described below.

### 2.2. Mass spectrometry

The bottom-up proteomics workflow, including sample preparation and liquid chromatography-tandem mass spectrometry (LC–MS/MS), closely followed our previously published reports [39, 40]. Briefly, 10 μg of recombinant repeat protein was precipitated in 70% acetone at –20 °C overnight and resuspended in 100 mM triethylammonium bicarbonate (TEAB) buffer with 0.2% deoxycholic acid (DCA), 6% sodium dodecyl sulfate (SDS), and 10 mM tris(2-carboxyethyl)phosphine (TCEP). Protein was denatured and reduced by incubation at 95°C for 10 min. Reduced protein was alkylated with 10 mM iodoacetamide (IAA) for 30 min at room temperature (RT) in the dark. The alkylation reaction was quenched by 1.2% phosphoric acid. The protein solution was spun onto an S-Trap device (Protifi, Farmingdale, NY) and digested by 750 ng trypsin (Promega, Madison, WI) in 100 mM TEAB buffer at 37°C overnight. Digested peptides were eluted by three buffers: 100 mM TEAB, 0.1% formic acid (FA) in water, and 50% acetonitrile (ACN) with 0.1% FA. Eluted peptides were combined and desalted using C18 ZipTips. Desalted peptides were reconstituted in water with 4% ACN containing 0.5% FA and analyzed by gradient elution–tandem mass spectrometry on a Waters NanoAcquity LC system coupled to a Q-Exactive mass spectrometer (Thermo).

### 2.3. Western blot

Recombinant repeats were resolved by 16% SDS-polyacrylamide gel electrophoresis (SDS-PAGE) and transferred to a polyvinylidene difluoride membrane. The membrane was blocked with 5% non-fat milk in TBS-T buffer (Tris buffered saline with 0.05% Tween 20) at RT for 1 hour. The membrane was hybridized at 4°C overnight with one of five primary antibodies: anti-6xHis (Thermo, HIS.H8, 1:2000), anti-CA125 epitope group A (Sigma, OC125, 1:200), anti-CA125 epitope group A (Fitzgerald, OC125-like, M61704, 1:2000), anti-CA125 epitope group B (Agilent, M11, 1:100), or anti-CA125 epitope group B (Fitzgerald, M11-like, M61703, 1:2000). The membrane was washed and hybridized with goat anti-mouse secondary antibody conjugated with horseradish peroxidase (HRP) (Thermo, 62-6520, 1:5000) at RT for 1 hour. Signals were developed using Pierce ECL Western Blotting Substrate (Thermo), and membranes were imaged on a ChemiDoc system (Bio-Rad, Hercules, CA).

### 2.4. ELISA

Pierce Nickel-Coated Plates (Thermo) were used to immobilize recombinant repeats through their 6xHis tag for indirect ELISA. Five hundred ng of recombinant repeat in 100 μL PBS buffer were used for immobilization. HE4 protein was used in place of recombinant repeats as a negative control, and wells with no immobilized protein were used as additional negative controls. After three washes with 200 μL PBS containing 0.01% Tween 20, 100 μL primary antibody—anti-6xHis (Thermo, HIS.H8, 1:2000), anti-CA125 epitope group A (Sigma, OC125, 1:200 or Fitzgerald, M61704, 1:2000) or anti-CA125 epitope group B (Agilent, M11, 1:100 or Fitzgerald, M61703, 1:2000)—was added to the well and incubated at RT for 1 hour. After three washes with 200 μL PBS-T, 100 μL anti-mouse HRP (Thermo, 62-6520, 1:2000) secondary antibody was added to the well and incubated at RT for 1 hour. Signals were developed using SuperSignal ELISA Femto Substrate (Thermo). Relative luminescence units at 425 nm were detected on a SpectraMax M5 plate reader (Molecular Devices, San Jose, CA).

### 2.5. Surface plasmon resonance

All SPR experiments were performed on a Nicoya Lifesciences OpenSPR-XT 2-channel instrument (Kitchener, ON) using Nicoya NTA-functionalized standard sensor chips. PBS with 0.05% Tween 20 (PBS-T; PBS from GrowCells, Irvine, CA; Tween 20 from Fisher Scientific, Waltham, MA) was used as both the immobilization and running buffer. Autosampler tray and sensor temperatures were set to 20°C. Recombinant repeat concentration was estimated on a ThermoFisher NanoDrop 2000 (Waltham, MA) using the Protein A_280_ function with the 1 Abs = 1 mg/mL setting. Recombinant repeats were diluted to 125 nM in PBS-T except for repeat 6, which required 500 nM to achieve acceptable immobilization levels. Regeneration was performed at the beginning of a run and after every ligand-analyte pair was injected, using two injections of 10 mM Glycine-HCl (VWR LifeSciences, Radnor, PA) and one injection of 350 mM EDTA (TCI America, Portland, OR).

After surface regeneration with EDTA, the surface was activated with 40 mM NiCl_2_ (BeanTown Chemical, Hudson, NH) and recombinant repeat was injected to achieve ligand immobilization. Recombinant His-tagged streptavidin (Fitzgerald Industries, Acton, MA) at a concentration of 0.75 μM was used as a negative control molecule, and then antibody was injected as the analyte. M11 was injected at stock concentration, OC125 was diluted 1:200, and both M11-like and OC125-like were diluted 1:2000. The surface was then regenerated, and the repeat was re-immobilized for the next antibody. Three replicates were collected per recombinant repeat (R5 had an additional replicate). All injections were at a flow rate of 20 µL/min, except for Glycine-HCl (150 µL/min) and EDTA (100 µL/min). All injections had a dissociation time of 270 s. Corrected signals (reference channel subtracted from ligand-immobilized channel) were used to calculate average signals for antibody binding over the interval 500–525 seconds at the end of the dissociation phase using TraceDrawer Analysis 1.9.2 (Ridgeview Instruments, Uppsala, Sweden).

## 3 Results

### 3.1. Recombinant tandem repeat protein expression and confirmation

Recombinant repeats were expressed in *E. coli* and purified on Ni-NTA beads. Yield and purity of material covered from Ni-NTA beads were confirmed by visualization on polyacrylamide gel with Coomassie blue staining (SI Figure 3). All recombinant repeats were first analyzed by Western blot with anti-6xHis antibody to confirm expression (Figure 1). A single band at the expected molecular weight (∼ 19.4 kDa) was observed for R2, R5, R6, R7 and R25. A minor additional band was observed for R9, R11, R34 and R58. The minor band could result from incomplete denaturation during SDS-PAGE sample preparation, protein degradation, or post-translational modification. The first two possibilities were ruled out by using stronger denaturing conditions for sample preparation and increasing the concentration of protease inhibitor in the lysis buffer. The recognition of both major and minor bands by anti-CA125 antibodies (Figure 1) suggests they are the same protein with different post-translational modifications that do not affect antibody binding. The amino acid sequences of recombinant repeats were verified by mass spectrometry using a proteomic workflow our lab previously reported for MUC16 detection. All recombinant repeat proteins have at least 95% coverage by mass spectrometry (Table 1 and SI Figure 2) except R6, for which 67% coverage was observed.

**Table 1.**
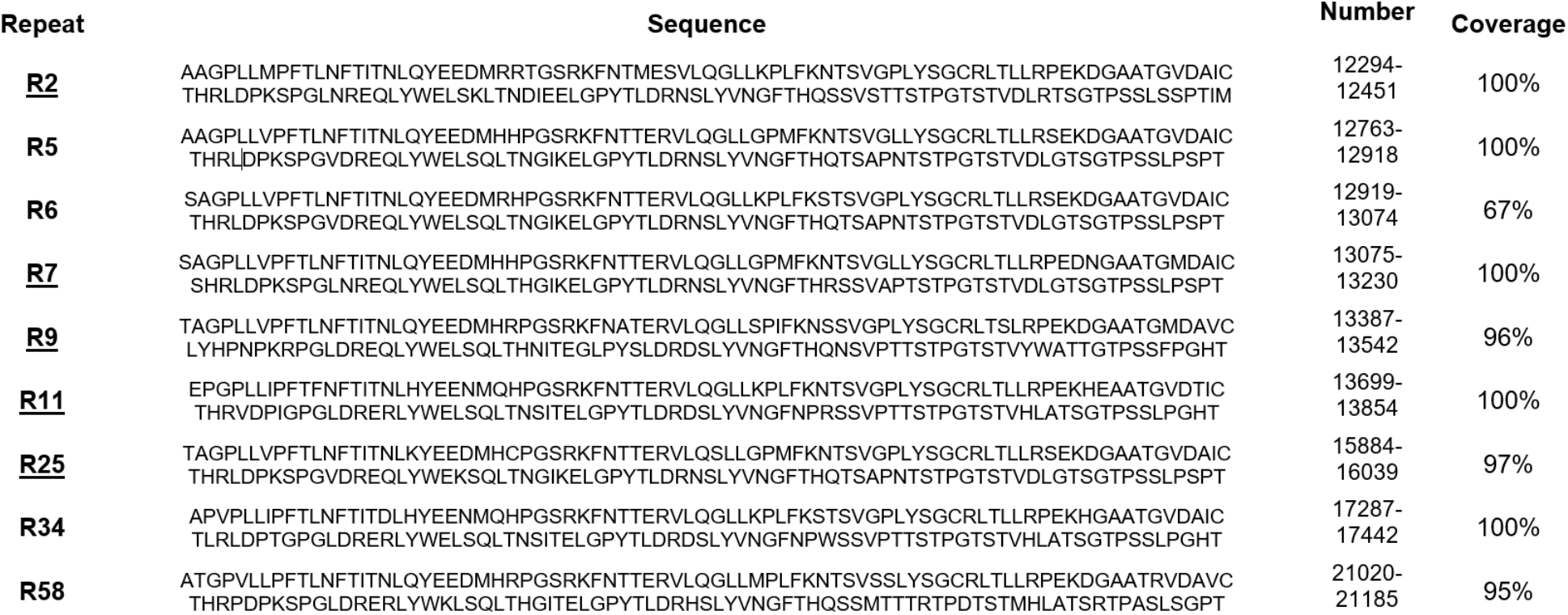
Recombinant repeats expressed and characterized in this study. Repeat numbering, amino acid sequences and corresponding amino acid number in reference sequence AF414442.2, as reported by O’Brien and co-workers [26], are shown in left three columns. The SEA domain extends from repeat 1 to 126 and is followed by a region rich in proline, serine, and threonine. Repeats with underlined numbers were previously examined by Bressan et al. [35]. Tandem repeat numbering increases from N- to C-terminus. Peptide coverage determined from bottom-up proteomics experiments to confirm expression is shown in the right column.

**Figure 2.**
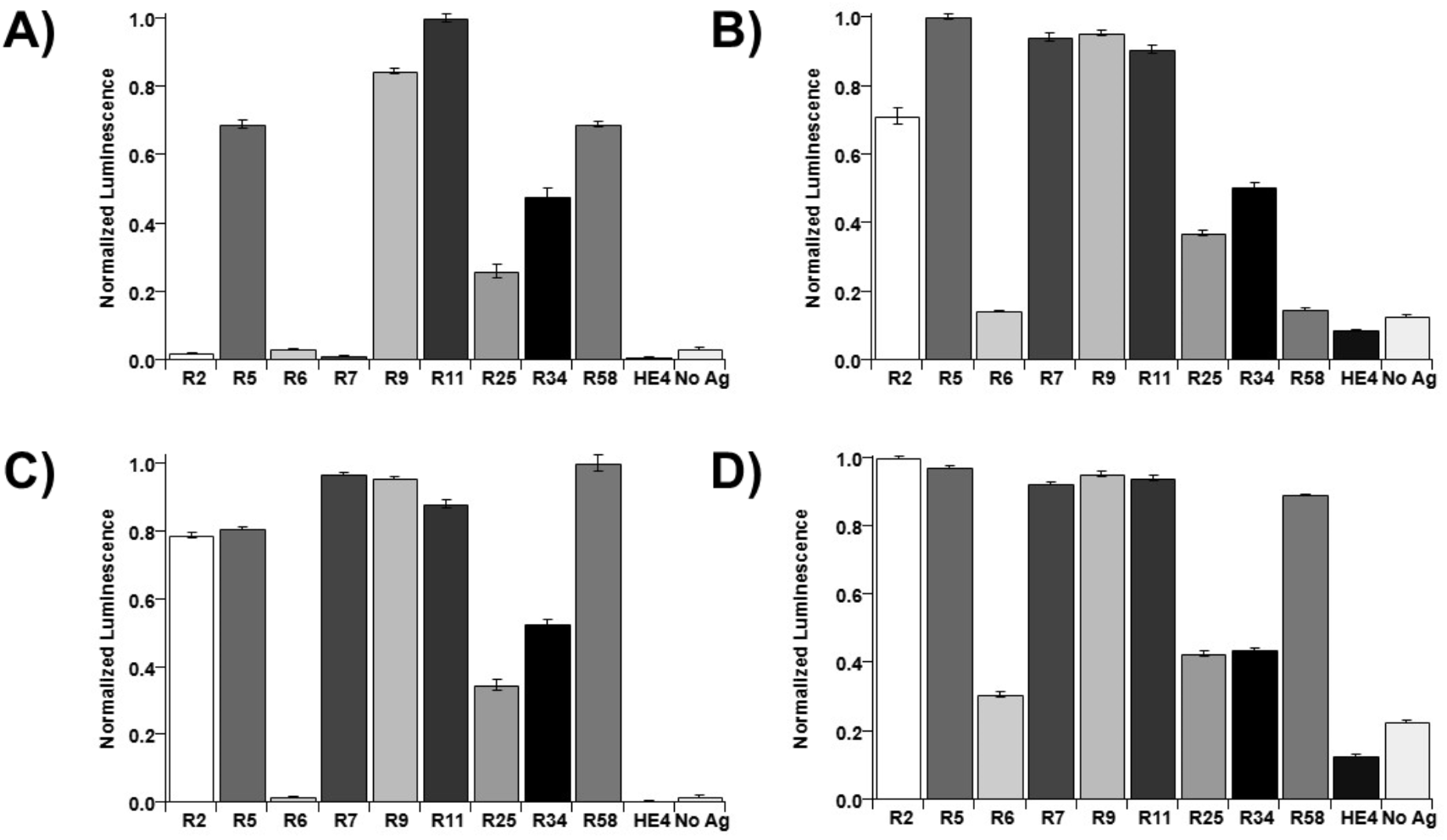
Results of indirect ELISA. Luminescent optical density values were normalized (to max = 1.0) within each data set. Recombinant repeats from MUC16 were probed with A) OC125 (1:200), B) M61704 (OC125-like, 1:2000), C) M11 (1:100), and D) M61703 (M11-like, 1:2000). Error bars are standard error of the mean (n = 3). HE4 and no antigen were used as negative controls.

**Figure 3.**
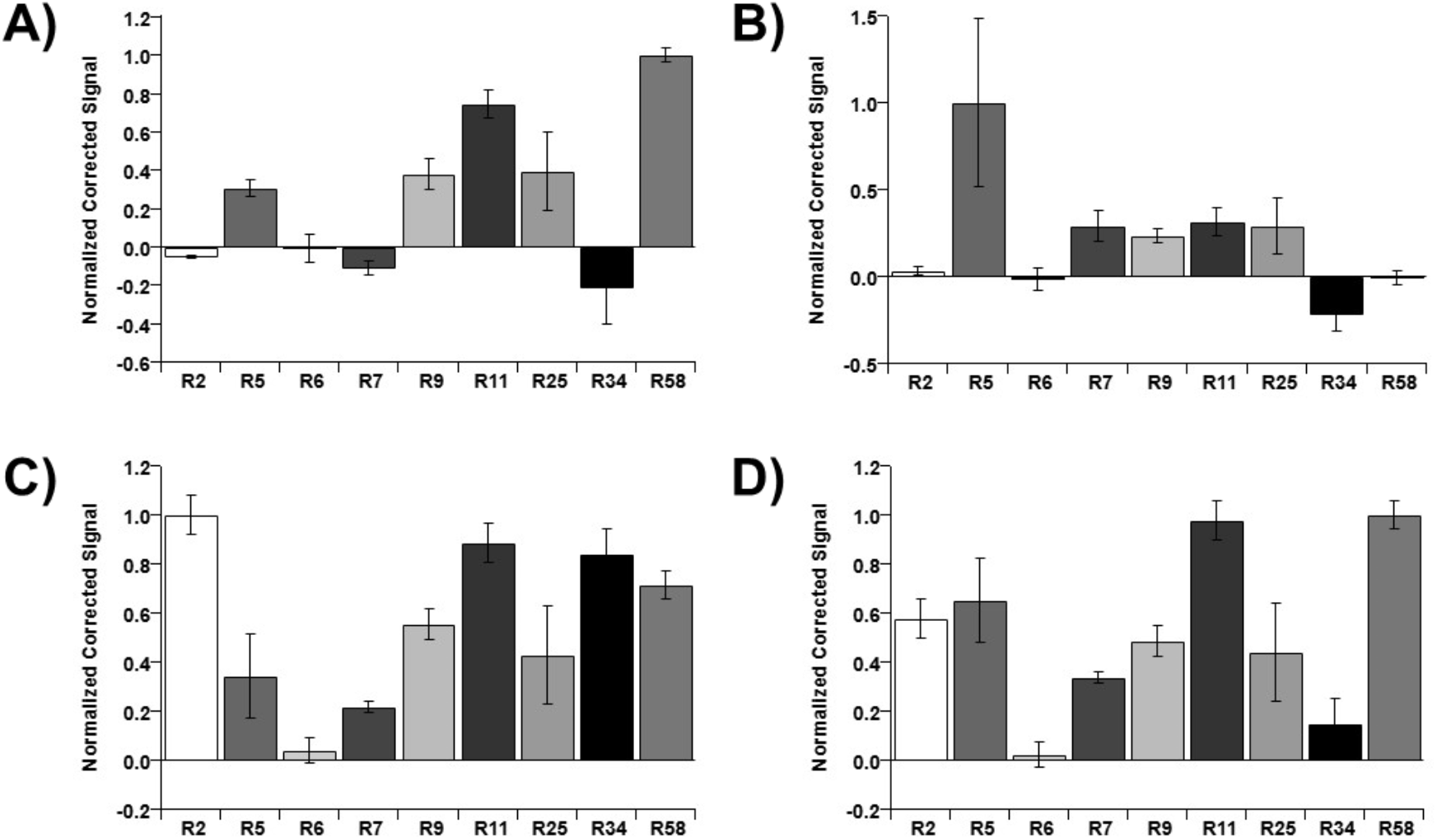
SPR data on binding of recombinant repeats from MUC16 to CA125 antibodies. Normalized (to max = 1.0) binding signals of A) OC125 (1:50), B) M61704 (OC125-like, 1:2000), C) M11 (stock concentration), and D) M61703 (M11-like, 1:2000) are shown. Error bars are standard error of the mean (n = 3, with the exception of R5, for which n = 4).

### 3.2. Western blot

Recombinant repeats were probed via Western blot with antibodies OC125, OC125-like, M11 and M11-like, and anti-6xHis antibody as a loading control (Figure 1). Epitope group A antibodies (OC125 and OC125-like), recognized repeats R5, R6, R9 and R25, with clear bands observed at the expected MW (∼19.4 kDa). Epitope group B antibody M11 recognized all repeats with different affinities, while M11-like antibody strongly bound to R2 and had faint bands for R5, R6 and R25. The uneven affinity among different recombinant repeats and antibodies is consistent with observations reported by Bressan and co-workers that epitopes were not uniformly distributed over the recombinant repeats they studied. In contrast to their observations, however, we do not detect binding between R11 and OC125 antibody, and we only observe weak binding of M11 antibody to R7 and R9. We observe that antibodies within the same epitope group have different recognition patterns at the level of individual expressed tandem repeats, and we suggest that this difference may also be observed on intact MUC16 *in vitro*. Further investigation is needed to confirm that these differences in binding are observed on intact MUC16.

### 3.3. ELISA

Recombinant repeats were immobilized onto Ni-NTA-coated 96-well plates for indirect ELISA. Luminescence values were normalized to the maximum signal for each antibody (Figure 2, shown as dot plots in SI Figure 4). Out of the four antibodies tested, OC125 showed evidence of binding to the fewest recombinant repeats (Figure 2A). OC125 showed maximum binding to R11 and strong binding to R5, R9, and R58, whereas R2, R6, and R7 had lower signals than the no-antigen control. OC125-like antibody (Figure 2B) displayed strong binding to R5, R7, R9, and R11, and little binding to R6 and R58. M11 (Figure 2C) had its maximum binding to R58, strong binding to R2, R5, R7, R9, and R11, and no apparent binding to R6. M11-like (Figure 2D) demonstrated maximum binding to R2, with strong binding to R5, R7, R9, R11, and R58. There were no repeats that showed no binding to any of the four antibodies, but R6 had smaller luminescence signals than the other recombinant repeats. This is not conclusive evidence that binding is not occurring, as the immobilization of protein within the wells was not quantified. The slightly higher luminescence signal in no antigen control wells compared to the HE4 controls indicates that nonspecific binding of antibodies to the wells may contribute to the luminescence signal. While the dilutions of antibodies varied, and the ranges of luminescence values varied from antibody to antibody, comparing the relative affinities of each clone to the repeats with one another can give some insight into the similarity of epitopes. Each antibody showed maximum binding signal to a different tandem repeat, indicating the binding locations may vary from antibody to antibody. OC125 and the OC125-like clone do not have similar patterns of binding from repeat to repeat (Figure 2A, 2B). M11 and the M11-like clone had similar binding patterns (Figure 2C, 2D).

**Figure 4.**
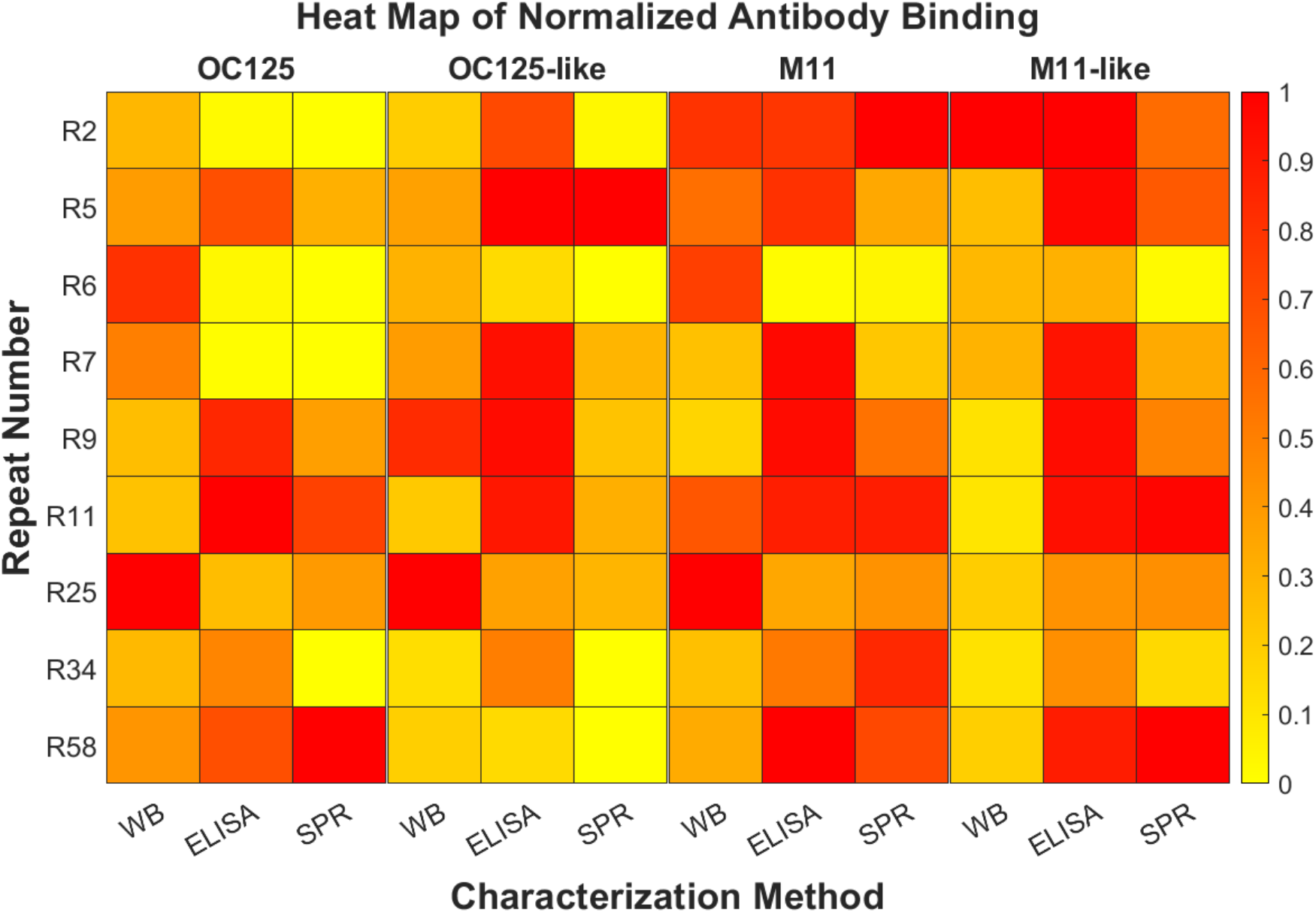
: Heat map generated using normalized binding signals of antibodies to recombinant repeats from MUC16. Data used to generate the heat map are in SI Table 1. Any negative normalized binding signals were set to zero for the generation of the heat map.

### 3.4. Surface plasmon resonance (SPR)

Representative sensorgrams are found in SI Figure 6. Average corrected binding signal for each antibody interacting with recombinant repeat was normalized to the maximum signal for each antibody (Figure 3, shown as dot plots in SI Figure 5). A negative normalized corrected value occurs when signal resulting from non-specific binding to the sensor chip is greater than signal from recombinant repeat–antibody binding. Due to variable antibody dilution, direct comparisons based on corrected signals cannot be made. However, the pattern of normalized binding for each recombinant repeat–antibody pair demonstrates variability across combinations of recombinant repeat and antibodies. OC125 (Figure 3A) and M11-like (Figure 3D) show the strongest response to R58, while M11 (Figure 3C) shows the strongest binding to R2, and OC125-like (Figure 3B) shows the strongest binding to R5. As with the Western blot and indirect ELISA, the differences in binding patterns between the clinical antibodies and their “-like” counterparts suggest that the location of their epitopes differs. OC125 and the OC125-like antibody show variability in binding patterns: OC125 shows the strongest binding response to R58, but OC125-like shows no binding to this repeat. Differences are also seen for M11 and the M11-like antibody: M11 shows stronger binding to R2 than to R5, while M11-like has similar affinity for both repeats. R34 and R58 also show different patterns of binding for M11 and M11-like: while the two repeats show comparable binding for M11, there is stronger binding to R58 for M11-like by a factor of 7.

## 4 Discussion

### 4.1. Comparison of analytical methods

Figure 4 displays the combined data for the three assays and enables visualization of differences in binding across recombinant repeat/antibody combinations (the raw data used to generate the heat map are found in SI Table 1). We observe that the extent of binding differs across the three analysis methods. One potential explanation of these differences is the nature of the assays: prior to Western blotting, recombinant repeats were denatured and treated to reduce disulfide bonds, while in ELISA and SPR assays the recombinant repeats remained as expressed with disulfide bonds intact. While it is not certain that the folded state of the expressed and purified recombinant repeats is structurally identical to the native state of a MUC16 tandem repeat *in vivo*, the discrepancy in antibody binding ability when recombinant protein is denatured supports the hypothesis that the epitopes for these antibodies is conformational [29, 36]. R6 and R25 are notable exceptions to the pattern. For these two recombinant repeats, Western blot results indicate antibody-repeat binding and the other binding analysis methods showed little evidence of interaction. In SPR experiments, R6 and R25 consistently had significantly lower immobilization signal response than other repeats, and yields during NTA affinity chromatography-based purification were consistently lower for these two recombinant proteins compared to the other seven. This observation implies that the lower binding signals in SPR and ELISA for R6 and R25 may result at least in part from inefficient immobilization of these recombinant repeats. The patterns of which repeat–antibody combinations show strong, weak, or minimal binding evidence look similar between ELISA and SPR, two methods in which the recombinant repeats were not denatured and were immobilized in the same orientation. ELISA generally showed more consistent binding signals across recombinant repeats than did SPR. Corrections for nonspecific binding could account for these differences. ELISA results may have been affected by variable immobilization or nonspecific binding, which were not tracked through data collection. In SPR assays nonspecific binding signal was subtracted from binding signal.

### 4.2. Working hypothesis regarding the nature of the CA125 epitopes

Structural models for each of the nine tandem repeat proteins were obtained using the iTasser server [41 – 43]. In all cases, the top threading template was 7sa9, the human MUC16 SEA5 domain reported by White et al. [23]. This structure was also the top identified structural analog in the Protein Data Bank for all nine tandem repeat proteins, with good topological similarity, inciated by TM-scores of 0.749, 0.760, 0.760, 0.761, 0.763, 0.761, 0.758, 0.765, and 0.758 for repeats R2, R5, R6, R7, R9, R11, R25, R34, and R58, respectively, where a TM score of 1 indicates a perfect match. The sequence identity of the tandem repeats with 7sa9 were 0.861, 0.967, 0.943, 0.885, 0.795, 0.811, 0.992, 0.811, and 0.836, indicating that this previously reported model is an excellent template for other MUC16 tandem repeat domains. The predicted models overlay the MUC16 SEA5 domains well (SI Figure 7), with the notable exception of the proline-serine-threonine (PST) rich region which is largely unstructured in the iTasser predictions of the tandem repeats. The SEA5 domain structure does not contain the PST region. The two major accessible faces noted by White et al. in their analysis of the crystal structure of SEA5 are seen in the tandem repeat domain proteins modeled here. It is not likely that the differences in antibody binding that we observe results from structural differences at these faces. Structural divergence between the SEA5 model (which contains a Serine at position 66) and the subset of tandem repeat proteins (R2, R7, R9, R11, R34, and R58) that contain Proline at that position is notable in the unstructured loop between beta strands that comprise the “C-loop” [32]. The center of the C-loop contains highly charged and polar amino acids and is found, in the model predicted by i-Tasser, to be adjacent to another loop that could be close enough to comprise a conformational epitope. Based on these studies, we hypothesize that the CA125 epitope involves the C-loop and charged and polar amino acids in the neighboring coil (residues 82-89). Notably, the tandem repeat protein that was found to be least immunogenic by Western blotting (R7) has aspartic acid replaced by asparagine within the loop that may be involved in conformational epitope presentation. Determining the exact location of the epitopes is the focus of our ongoing work.

## 5 Conclusions

This study contributes to ongoing efforts to identify the peptide epitopes of CA125. We anticipate that CA125 can be a more informative clinical biomarker if the nature of its recognition by antibodies is elucidated. Such knowledge may enable development of affinity reagents that recognize all repeats of MUC16. The current clinical test may under-report concentrations of CA125 in blood samples if the antibodies used for detection do not reliably recognize the subdomains in the immunogenic region, as data reported here suggest. Detection of smaller amounts of CA125 may be possible, enabling earlier detection of CA125 resurgence, which correlates strongly with cancer recurrence. Future work following from this study includes structural characterization of free and antibody-bound recombinant repeats and the use of recombinant repeats as targets for the generation of novel affinity reagents such as nucleic acid aptamers, immunoaffinity reagents, and vaccines.

## Acknowledgments

The authors thank Bill Boggess and the Notre Dame Mass Spectrometry and Proteomics Facility for technical assistance and Professor Brian Blagg for use of a ChemiDoc imager. The authors thank Professor Matthew Champion for helpful discussions and access to equipment for bacteria culture. Naviya Schuster-Little provided the structural alignment in SI Figure 1, and we thank her for many helpful discussions about protein structure prediction and comparisons. Daryl Good and Marko Jovic at Nicoya were pivotal in developing and optimizing SPR assays. This work was supported by award R21CA267532 from the National Cancer Institute and a Medical Research Program award from Tell Every Amazing Lady® About Ovarian Cancer Louisa M. McGregor Ovarian Cancer Foundation (T.E.A.L.®). Additional support came from a University of Notre Dame Advancing Our Vision Award in Analytical Science and Engineering (to RJW) and from the Department of Chemistry and the Office of Research at the University of Kansas.

## Author contributions

Conception: CWW, EKH, LM, RJW

Interpretation or analysis of data: CWW, EKH, LM, RJW

Preparation of the manuscript: CWW, EKH, LM, RJW

Revision for important intellectual content: CWW, EKH, LM, RJW

Supervision: RJW

---

**SI Figure 1:**
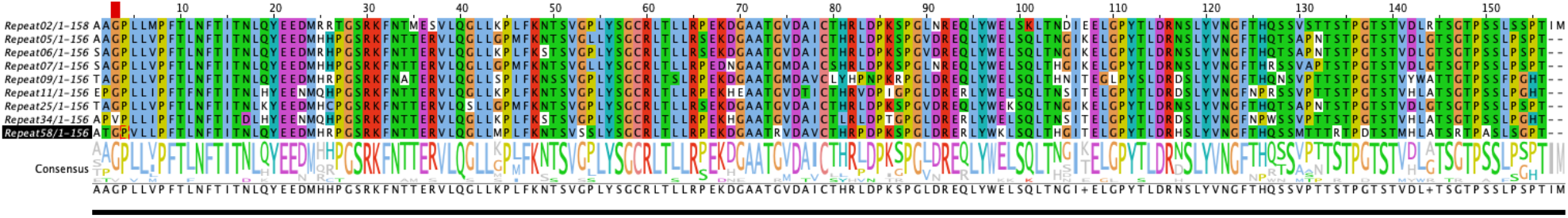
Sequence alignment of the MUC16 tandem repeat proteins. The alignment was created using Clustal Omega (https://www.ebi.ac.uk/Tools/msa/clustalo/) and visualized using Jalview.

**SI Figure 2:**
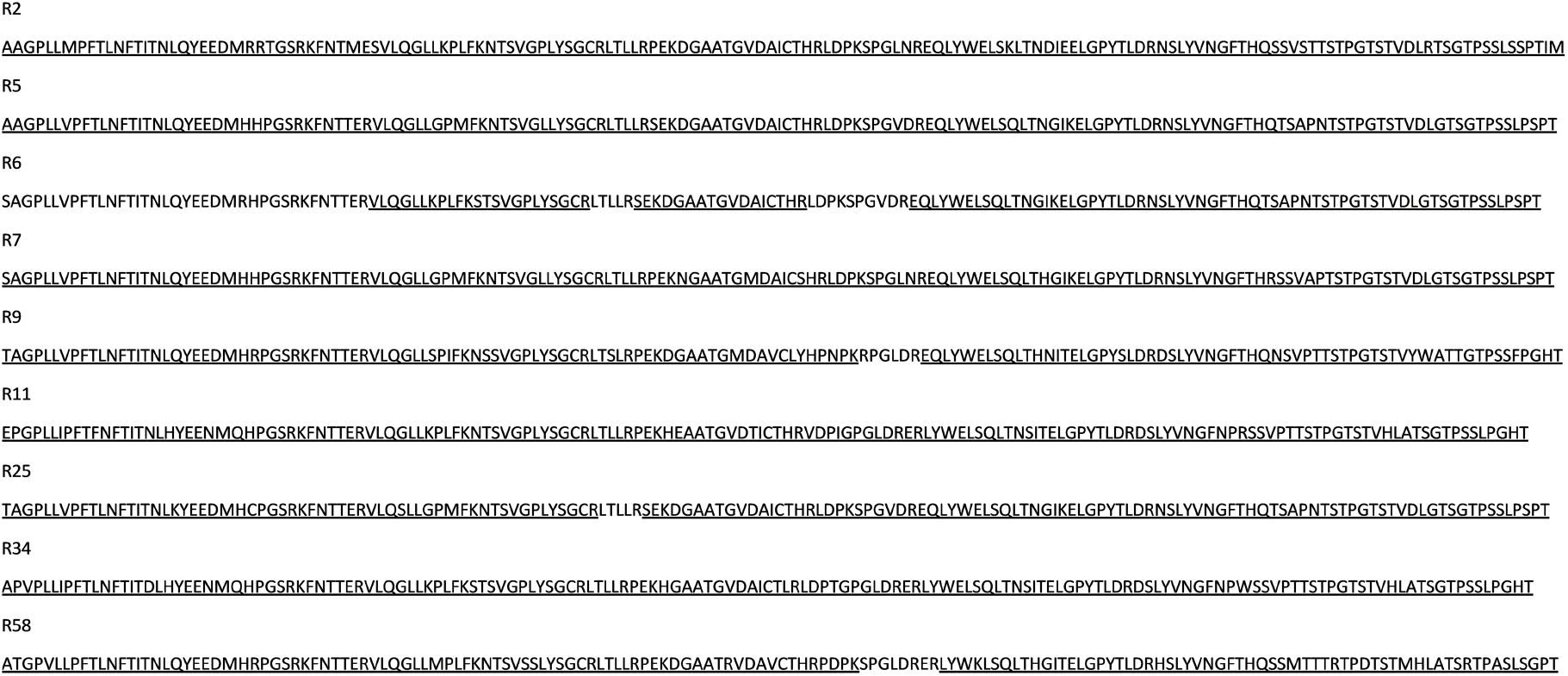
Mass spectrometry coverage map. The nine recombinant repeats studied here are shown in one-letter amino acid notation. Underlined amino acids were detected in tryptic peptides via LC-MS/MS.

**SI Figure 3.**
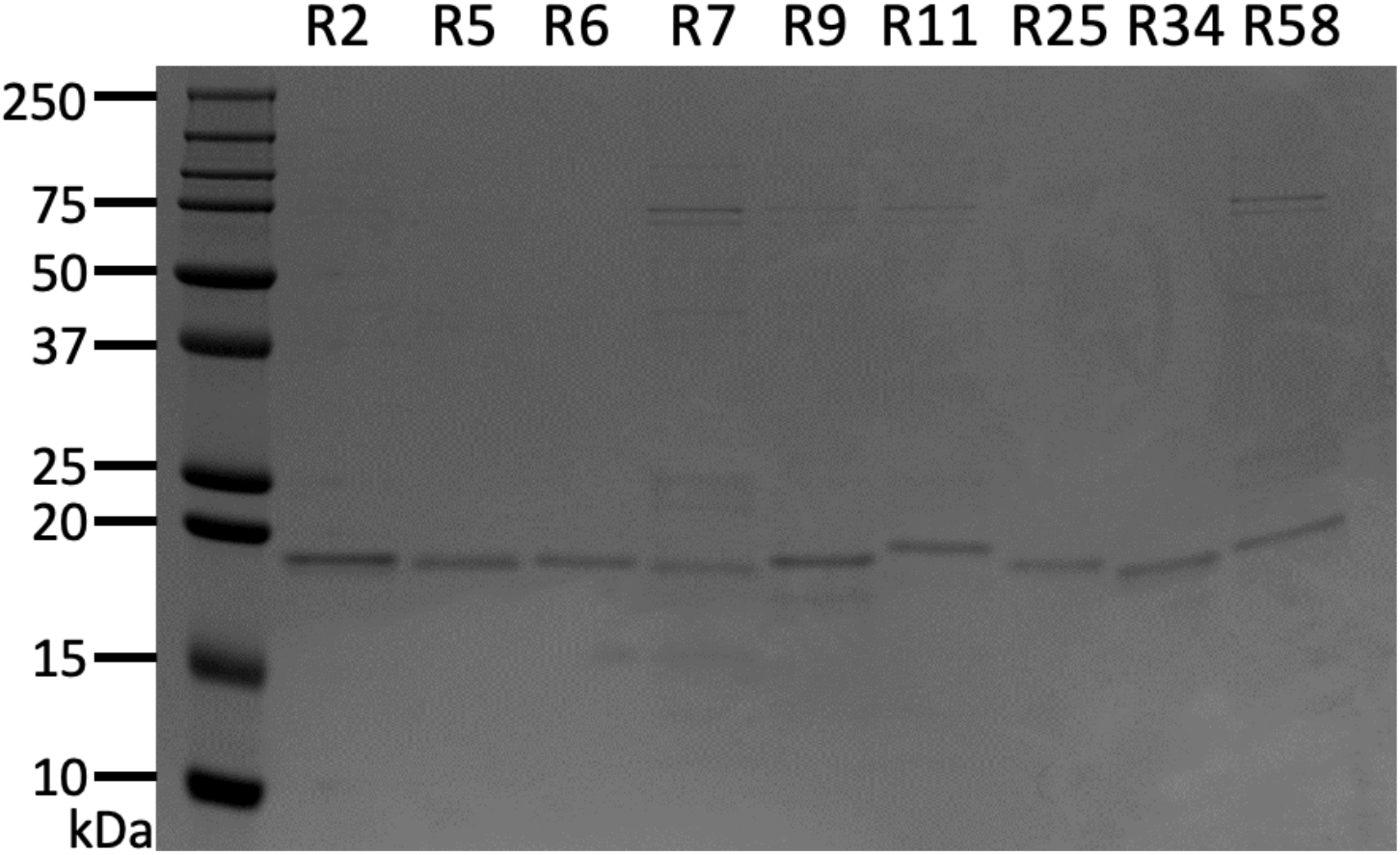
Coomassie blue stained gel image of recombinant repeats from MUC16 purified on Ni-NTA beads. Proteins eluted from Ni-NTA beads were separated by 16% SDS-PAGE at 100V for 2 hours. The gel was stained by Coomassie brilliant blue R-250 (Bio-Rad) and imaged on an Azure 400 imaging system (Azure).

**SI Figure 4.**
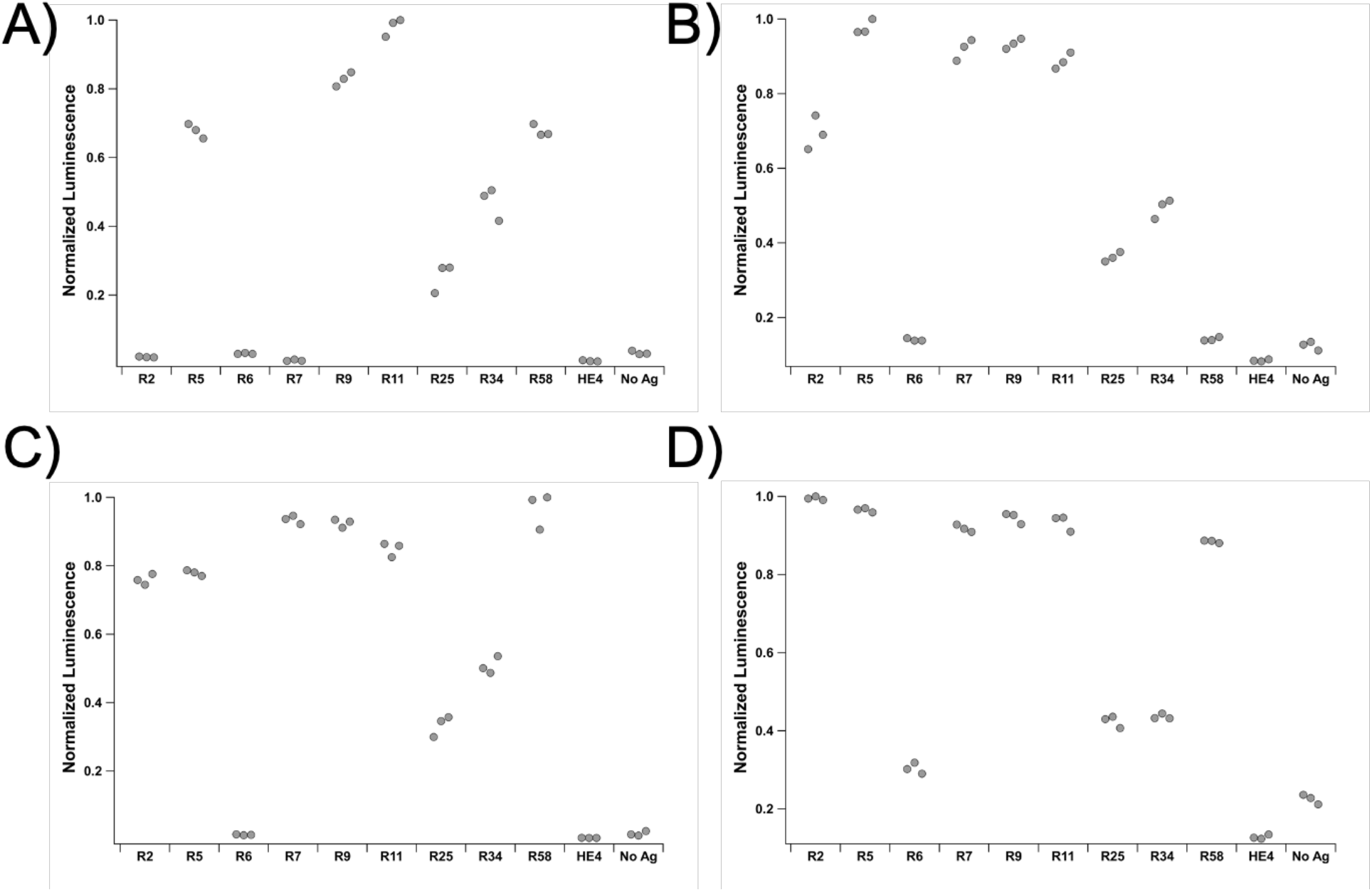
Results of indirect ELISA, same as Figure 2, shown as a dot plot. Luminescent optical density values were normalized (to max = 1.0) within each data set. Recombinant repeats from MUC16 were probed with A) OC125 (1:200), B) M61704 (OC125-like, 1:2000), C) M11 (1:100), and D) M61703 (M11-like, 1:2000).

**SI Figure 5.**
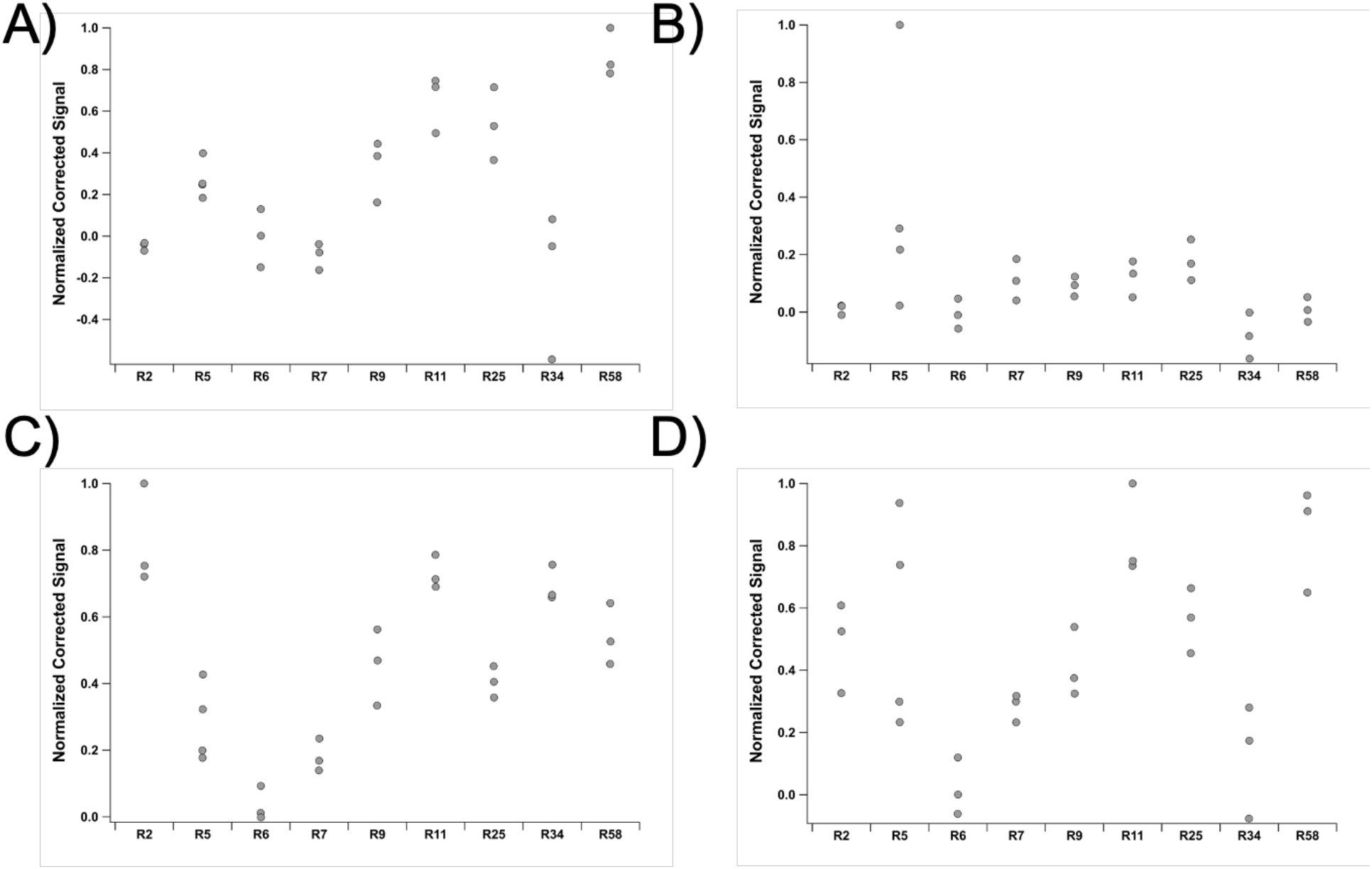
SPR data on binding of recombinant repeats from MUC16 to CA125 antibodies, same as Figure 3, shown as a dot plot. Normalized (to max = 1.0) binding signals of A) OC125 (1:50), B) M61704 (OC125-like, 1:2000), C) M11 (stock concentration), and D) M61703 (M11-like, 1:2000) are shown.

**SI Figure 6:**
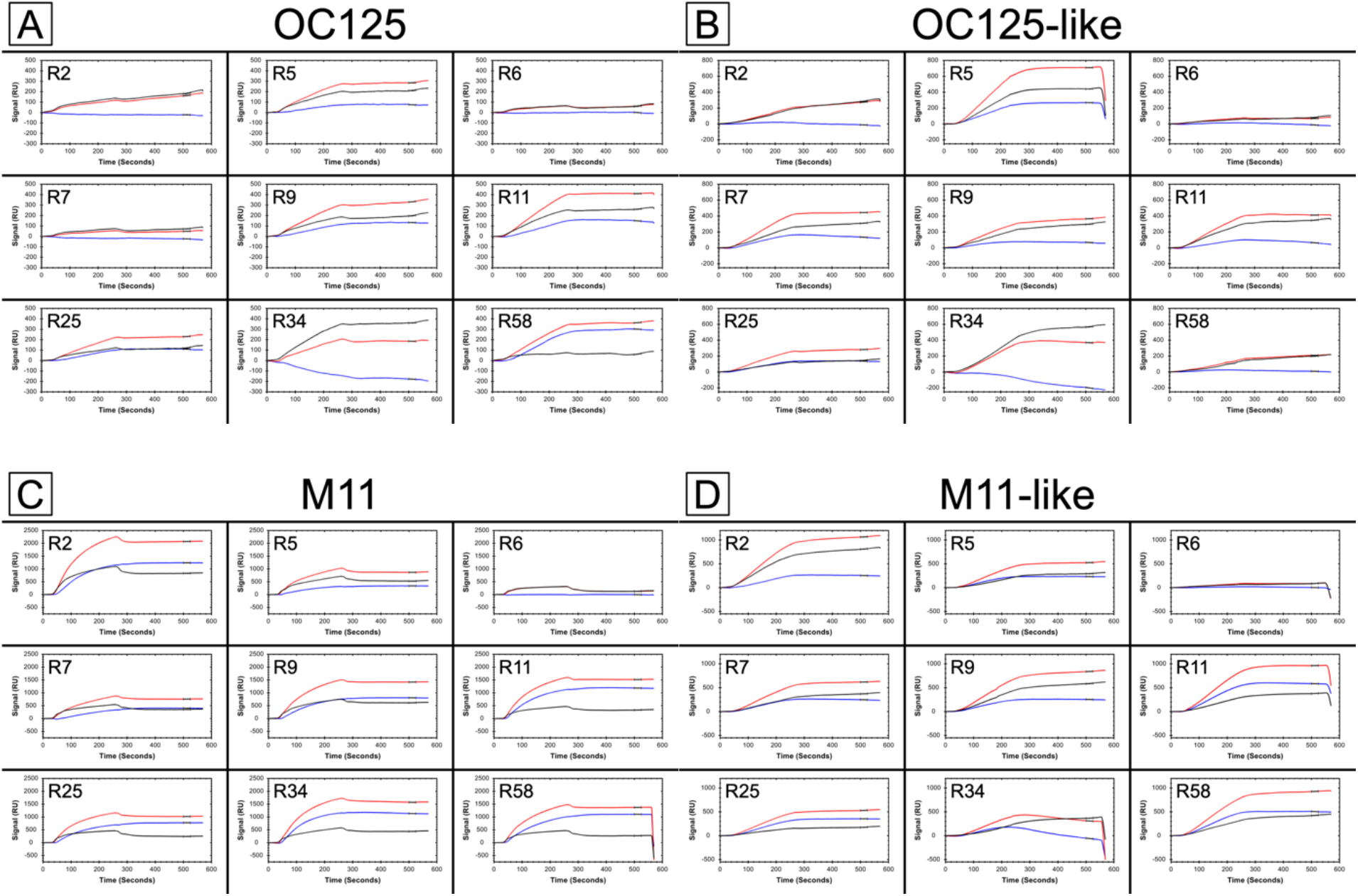
Representative sensorgrams displaying real-time binding of antibodies to recombinant repeats. A) OC125, B) OC125-like, C) M11, and D) M11-like. The black trace corresponds to Channel 1 signal, the red trace corresponds to Channel 2, and the blue trace shows Corrected Signal (Channel 2 – Channel 1). The average value of the Corrected Signal over the interval between 500 and 525 seconds was used for quantitative assessment of binding.

**SI Figure 7:**
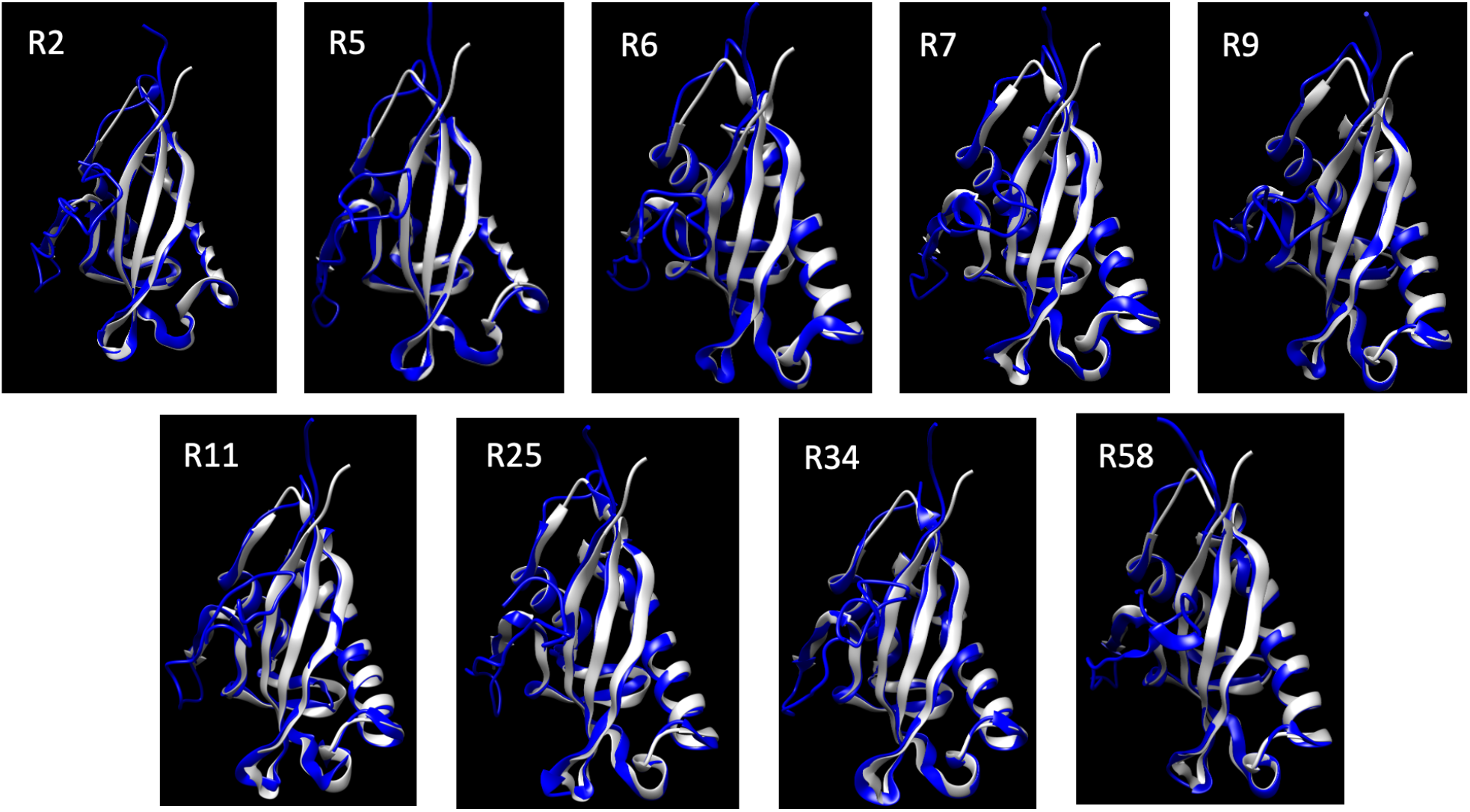
Ribbon models of the MUC16 tandem repeat proteins predicted by iTasser (blue) and overlaid with the human SEA domain 5 (7sa9) as determined by White et al. (2022) (light grey). The cysteine loop (C-loop) comprises the middle of the right two antiparallel beta strands and the intervening loop (oriented toward the bottom of the image). Overlaid plots prepared in Chimera.

**SI Table 1:** Data used to generate heat map in Figure 4.

|  | **OC125** |  |  | **OC125-like** |  |  | **M11** |  |  | **M11-like** |  |  |
| --- | --- | --- | --- | --- | --- | --- | --- | --- | --- | --- | --- | --- |
|  | **WB** | **ELISA** | **SPR** | **WB** | **ELISA** | **SPR** | **WB** | **ELISA** | **SPR** | **WB** | **ELISA** | **SPR** |
| **R2** | 0.283 | 0.020 | -0.054 | 0.197 | 0.711 | 0.029 | 0.797 | 0.786 | 1 | 1 | 1 | 0.575 |
| **R5** | 0.388 | 0.691 | 0.31 | 0.369 | 1 | 1 | 0.565 | 0.807 | 0.342 | 0.255 | 0.970 | 0.652 |
| **R6** | 0.805 | 0.030 | -0.007 | 0.302 | 0.144 | -0.018 | 0.752 | 0.014 | 0.042 | 0.279 | 0.305 | 0.023 |
| **R7** | 0.502 | 0.010 | -0.107 | 0.389 | 0.941 | 0.291 | 0.246 | 0.968 | 0.219 | 0.297 | 0.923 | 0.334 |
| **R9** | 0.257 | 0.844 | 0.378 | 0.830 | 0.956 | 0.237 | 0.165 | 0.957 | 0.552 | 0.114 | 0.950 | 0.488 |
| **R11** | 0.239 | 1 | 0.746 | 0.211 | 0.908 | 0.316 | 0.661 | 0.879 | 0.885 | 0.103 | 0.938 | 0.978 |
| **R25** | 1 | 0.26 | 0.395 | 1 | 0.371 | 0.291 | 1 | 0.346 | 0.426 | 0.198 | 0.426 | 0.440 |
| **R34** | 0.274 | 0.479 | -0.213 | 0.130 | 0.505 | -0.216 | 0.248 | 0.526 | 0.841 | 0.117 | 0.438 | 0.149 |
| **R58** | 0.417 | 0.691 | 1 | 0.188 | 0.145 | -0.010 | 0.329 | 1 | 0.715 | 0.188 | 0.889 | 1 |

